# Early subcortical response at the fundamental frequency of continuous speech measured with MEG

**DOI:** 10.1101/2023.06.23.546296

**Authors:** Alina Schüller, Achim Schilling, Patrick Krauss, Tobias Reichenbach

## Abstract

Most parts of speech are voiced, exhibiting a degree of periodicity with a fundamental frequency and many higher harmonics. Some neural populations respond to this temporal fine structure, in particular at the fundamental frequency. This frequency-following response to speech (speech-FFR) consists of both subcortical and cortical contributions and can be measured through electroencephalography (EEG) as well as through magnetoencephalography (MEG), although both differ in the aspects of neural activity that they capture: EEG is sensitive to both radial and tangential sources as well as to deep sources, while MEG is more restrained to the measurement of tangential and superficial neural activity. EEG responses to continuous speech have shown an early subcortical contribution, at a latency of around 9 ms, in agreement with MEG measurements in response to short speech tokens, whereas MEG responses to continuous speech have not yet revealed such an early component. Here we analyze MEG responses to long segments of continuous speech. We find an early subcortical response at a latency of 9 ms, followed by later right-lateralized cortical activities at delays of 20 - 57 ms as well as potential subcortical activities. Our results show that the early subcortical component of the FFR to continuous speech can be measured from MEG, and that its latency agrees with that measured with EEG. They furthermore show that the early subcortical component is temporally well separated from later cortical contributions, enabling an independent assessment of both components towards further aspects of speech processing.

## INTRODUCTION

Speech is a highly complex acoustic signal that needs to be processed in the brain in real time for comprehension. Investigations of the neural mechanisms that yield such rapid processing are increasingly employing more natural stimuli, from individual syllables and words to sentences and even entire stories [11, 34]. Such studies have, for instance, revealed cortical tracking of characteristic, slow rhythms in speech set by the rates of phonemes, syllables and words [11, 20, 21, 25, 59].

Faster neural activity reflects the temporal fine structure of voiced speech such as vowels or voiced consonants. During the production of these speech parts, the vocal chords vibrate at a certain fundamental frequency *f*_0_, typically between 100 - 300 Hz, resulting in a periodic signal [6]. The fundamental frequency and its higher harmonics constitute the signal’s temporal fine structure [23, 53].

A subcortical response to the temporal fine structure of speech can be measured noninvasively in humans using electroencephalography (EEG) as well as magnetoencephalography (MEG) [8, 14, 16, 29]. Moreover, such measurements of the frequency-following response to speech (speech-FFR) through EEG or MEG with source reconstruction have recently identified contributions from the auditory cortex [8, 13–15, 32, 43].

While the spatial origins of the different neural contributions to the speech-FFR have been increasingly clarified, the temporal aspects remain less clear. Neural activity in the inferior colliculus is assumed to occur at a delay of 5 - 7 ms [48]. However, EEG measurements of FFRs often find somewhat longer latencies of around 9 ms and up to 14 ms [7, 27, 39, 40]. MEG measurements of cortical contributions to the speech-FFR have an even larger uncertainty around the timing, pinning the response between 12 - 60 ms [13, 43]. Because the earliest sound-evoked neuronal activities in auditory cortex can occur already 9 ms after a stimulus onset, the cortical contributions to the speech-FFR might overlap in time with subcortical contributions.

A better understanding of the timing and source of the different components of the speech-FFR matters for investigating the role of these different components for speech processing. The speech-FFR was found to be related to frequency discrimination, to language experience, and to musical expertise [41, 42, 47, 49, 60]. Moreover, we demonstrated that the EEG-measured response is modulated by selective attention to one of two competing speakers, presumably due to top-down feedback from higher cortical areas [24, 27]. However, a more precise understanding of the involved neural feedback loops requires a better spatiotemporal segregation of the different neural activities.

MEG and EEG measurements of the speech-FFR have predominantly employed short speech tokens such as single vowels or syllables and achieved high accuracy of spatial source localization due to the use of a high number of repetitions [8, 13–15]. However, the temporal spread in the autocorrelation of the voiced parts of these short speech signals limited the temporal resolution of the neural activities.

In particular, a recent MEG study estimated the response delay to a single repeated syllable through computing the explanatory power over successive windows of 12 ms in duration [14]. This power was found to increase between 0 ms and 36 ms for the subcortical structures, and between 0 ms and 48 ms for the auditory cortex. This comparatively coarse estimate of the different delays left unclear to which degree subcortical and cortical activities might temporally overlap. A recent EEG investigation into the speech-FFR elicited by repeated presentation of a short speech token did not determine the delays of the responses from the subcortical and cortical sources, but their relative delays, obtaining significant spread in these latencies [8]. Another recent MEG experiment investigated the neural responses to pure tones of different frequency [29]. Although it could successfully discriminate between different subcortical and cortical contributions to the FFR, it did not allow to obtain timing estimates of the different sources.

As another approach, we recently employed continuous speech to measure the speech-FFR with EEG [24, 27]. We therefore extracted a fundamental waveform from the speech signal that, at each time instance, oscillates at *f*_0_. This waveform could then be related to the EEG recordings through regularized regression, yielding temporal response functions (TRFs) that show the contribution of different scalp electrodes at different delays. This statistical approach allowed for an estimation of the delay of the response at about 9 ms, indicating that only subcortical contributions were measured.

A similar study analyzed MEG responses to continuous speech and found neural sources between 23 - 63 ms, indicating that mostly cortical contributions were recorded [43]. The lack of subcortical activity was presumably due to the low sensitivity of MEG for deeper sources and the relatively short speech material of six minutes per subject.

Here we sought to measure and spatiotemporally localize both subcortical and cortical contributions to the speech-FFR using continuous speech as measured from MEG. To compensate for the poor sensitivity of MEG to subcortical activity, we employed comparatively long MEG measurements of 17 minutes per subject (Fig. 1). We then performed a spatio-temporal source reconstruction to differentiate and locate the neural activities.

**FIG. 1.**
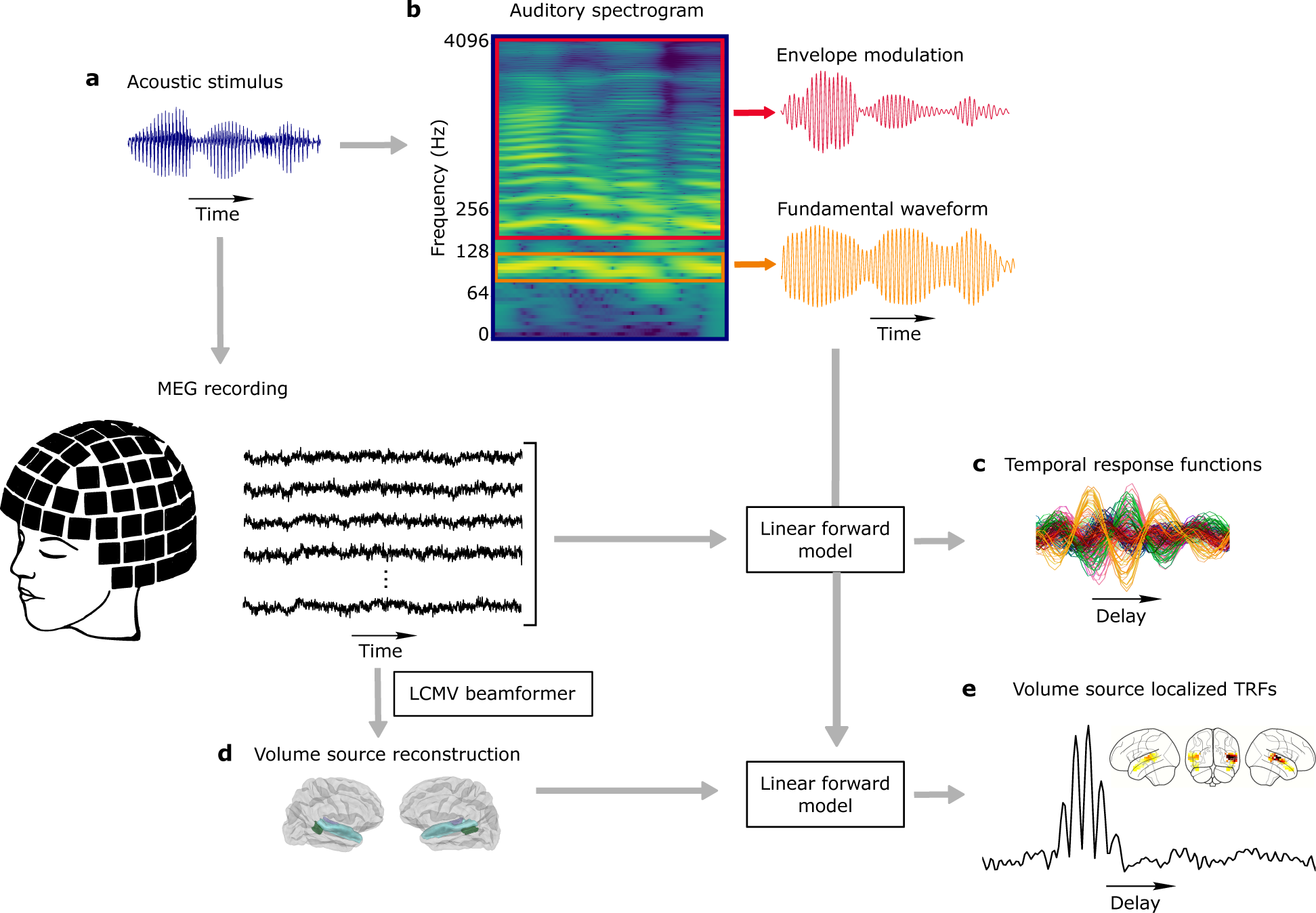
Overview of the experimental setup and data analysis. (a) We measured MEG (black) in response to continuous speech (blue). (b) A spectrogram was computed from the audio input to extract the fundamental waveform (orange) and the envelope modulation (red). (c) Sensor-level Temporal Response Functions (TRFs) were calculated for both audio features through a linear forward model, which estimates the neural response from the speech features. (d) Volume source reconstruction was performed on an average MRI brain template for two regions of interest (ROIs) by applying a linearly constrained minimum variance (LCMV) beamformer to the raw MEG data. (e) Volume source localized, i.e. source-level, TRFs were calculated for both audio features through the linear forward model.

## MATERIALS AND METHODS

### Experimental Design

We employed an existing experimental dataset that was collected for the study of continuous neuronal activity evoked by natural speech [55]. MEG recordings were obtained from 15 healthy right-handed monolingual native German speakers (20-42 years, 8 females, 7 males).

Participants had no history of neurological illness, drug abuse or hearing impairment. The study was granted ethical permission by the ethics board of the University Hospital Erlangen. The number of participants was chosen based on previous studies on neural responses to speech [11, 24, 43].

The participants listened to continuous speech, presented in form of an audio book, which was based on the German novel *Gut gegen Nordwind*, written by Daniel Glattauer and published by *Hörbuch Hamburg*. The audio book is available in stores and permission to use it for current and future studies has been granted by the publisher. The audio book has a total duration of 4.5 h. One female and one male speaker narrate alternately without a competing talker or other background noise. The auditory stimuli were presented at approximately 50 dB SPL. Small adjustments to the sound-pressure level were carried out for a few participants to ensure they could hear the sound comfortably.

The first 40 minutes of the audio book were presented diotically to the participants in ten parts. After each part, three multiple-choice comprehension questions had to be answered on a monitor, to test attention. The MEG recording during these breaks was eliminated from the analysis. Furthermore, the acoustic stimulation was stopped two times for a five-minute break. This resulted in a total experimental duration of approximately one hour. For this study, we only considered the MEG responses to the male speaker (17 min of the 40 min audio stimulus) due to its lower fundamental frequency, leading to larger neural responses [57].

MEG data (248 magnetometer, 4D Neuroimaging, San Diego, CA, USA) were recorded with a sampling frequency of 1, 017.25 Hz (supine position, eyes open, analogue band-pass filtering between 0.1 *−* 200 Hz). By the use of an integrated digitizer (Polhemus, Colchester, Vermont, Canada), five landmark positions were recorded and a calibrated linear weighting of 23 reference sensors (manufacturing algorithm, 4D Neuroimaging, San Diego, CA, USA) was used to correct for environmental noise. The collected data was further processed by applying a digital band-pass filter (70 *−* 130 Hz) offline for speech feature analysis, as well as a 50 Hz notch filter. The data was furthermore downsampled to a sampling frequency of 1, 000 Hz.

The speech signal was presented simultaneously to the MEG recording (Fig. 1 (a), (b)) through a custom-made setup that is described in detail by Schilling et al. [55]. A stimulation computer was connected to an external USB sound device, which provided five analogue outputs. The first two of these outputs were connected to an audio amplifier, of which the first output was connected in parallel to an analogue input channel of the MEG data logger. An alignment of the speech stimulus to the MEG recording with an accuracy of 1 ms could be achieved through cross-correlating the speech stimulus with the audio reference recording obtained by the analogue input channel of the MEG data logger.

### Data Analysis

#### Acoustic stimulus representations

We used two speech features to investigate the speech-FFR from MEG recordings, the fundamental waveform as well as the high-mode envelope modulation (Fig. 1 (b)).

The first feature, the fundamental waveform, was computed through applying a bandpass-filtering to the speech signal between 70*−*130 Hz, that is, around the fundamental frequency that was, on average 95 Hz. The so-obtained fundamental waveform is very similar to that obtained from empirical mode decomposition [24, 27, 36]. The neural response to the fundamental waveform captures the neural activity that emerges directly in response to the fundamental frequency, and is sometimes referred to as spectral FFR [1].

Regarding the second feature, previous studies showed that the neural response at the fundamental frequency of speech is also driven by the envelope modulation of the higher harmonics [38, 43]. We therefore computed the envelope modulation by first extracting frequency bins, each of size 100 Hz, in the range from 250 Hz to 4, 050 Hz using band-pass filters. This resulted in 38 extracted higher modes of which the acoustic envelopes were calculated. All higher-mode envelopes were then band-pass filtered between 70 *−* 130 Hz, that is, in the same range as the fundamental waveform. This procedure resulted in a band-limited envelope for each frequency bin. Averaging across the envelopes yielded the high-mode envelope modulation that contained the acoustic information of all frequencies up to 4, 050 Hz. The corresponding neural response has previously also been referred to as envelope FFR [1].

The applied band-pass filters were implemented using Python’s *Scipy* library [58]. A linear digital Butterworth filter (second order, critical frequencies obtained by dividing the lower and upper cut-off frequency by the Nyquist frequency) was applied twice, once forward and once backwards, to prevent phase delays.

#### Temporal Response Functions (TRFs)

To investigate the origin of the neural response to continuous speech measured with MEG, we computed TRFs for the MEG channels as well as for the estimated vertices in the source space. We therefore applied a linear forward model that reconstructed the multi-channel MEG response *y*^(^*^c^*^)^ at each MEG channel (or source voxel) *c* and at time *t* from a linear combination of acoustic stimulus samples, shifted by time delays *τ* that ranged from a minimal value *τ*_min_ to a maximal value *τ*_max_:

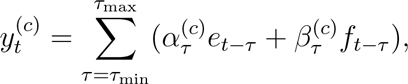

where *e_t−τ_* and *f_t−τ_* describe the time-delayed envelope modulation and fundamental waveform, respectively. The weights *α*^(^*^c^*^)^ and *β*^(^*^c^*^)^ of this linear combination are referred to as the temporal response functions (TRFs) of the two acoustic stimulus features (Fig. 1 (c)). A TRF can be viewed as the set of weights that best describe the time course of the neural response *y*^(^*^c^*^)^ to a feature at each channel (or source voxel) *c* and therefore gives rise to the neural response to each acoustic feature across the different time lags *τ*.

The TRFs were computed for time lags ranging from *τ*_min_ = *−*20 ms to *τ*_max_ = 140 ms, with an increment of 1 ms, corresponding to a sampling frequency of 1, 000 Hz. This resulted in 161 time lags in total. Although we did not expect any neural response to occur at negative time lags, where the acoustic stimulus lagged behind the neural response, or for time lags larger than 100 ms, we still took both temporal ranges into account to control for the absence of significant responses there.

The TRF coefficients were estimated using regularized ridge regression [33]. The regularization parameter *λ* can thereby be expressed as *λ* = *λ_n_ · e_m_*, in which *e_m_* is the mean eigenvalue of the covariance matrix, and *λ_n_* is the normalized regularization parameter. For the TRF estimations in this study, we used a fixed normalized regularization parameter *λ_n_* = 0.1 across all subjects. This value was chosen based on cross-validation, which was applied to all subjects individually and yielded an optimum around *λ_n_* = 0.1 for each subject. The implementation of the forward model and the TRF estimation employed the algorithms developed by Etard et al. [24] and Kegler et al. [38].

Due to an incomplete dataset for two of the 15 participants, we excluded those from the further analysis. The TRFs were estimated for each of the 13 subjects (20-42 years, 7 females, 6 males) individually. The results were then averaged across subjects, yielding population average models. To obtain one TRF magnitude value for each time lag, the average across MEG channels (or across vertices) was taken.

#### Neural source estimation

The neural sources of the MEG signals were computed using the MNE-Python software package [30]. Since no subject-specific MR-scans were available, we used the Freesurfer template MRI ‘fsaverage’ [26]. It is worth noting that the use of an average brain template can provide comparable results to individual MR scans in source localization analyses [22, 35] and has been validated in previous research on neural mechanisms of speech processing [43]. The head position of each subject with respect to the MEG scanner was recorded in the beginning and at the end of each measurement with five marker coils. Moreover, the head shape was digitized (Polhemus, Colchester, Vermont, Canada). The so obtained subject-individual information were used to coregister the ‘fsaverage’ brain template using rotation, translation and uniform scaling. For one subject, there was no head digitization recorded and we therefore excluded this participant from the source reconstruction analysis, in addition to the two prior excluded subjects, resulting in twelve subjects (20-42 years, 6 females, 6 males) for the source analysis.

We created a volumetric source space for the average brain. The volume source space was defined on a regular grid of 5 mm spacing between neighboring grid points and the Freesurfer ‘aparc+aseg’ parcellation was applied to define regions on which the sources were estimated. We then divided the volume source space into a cortical and a subcortical portion. The cortical space included the auditory cortex on both the right and left hemisphere (‘aparc’ labels: ‘transversetemporal’, ‘superiortemporal’ and ‘bankssts’), leading to 202 source locations with arbitrary orientations. The subcortical region contained the brainstem (‘aseg’ labels: ‘Brain-Stem’), resulting in 207 source locations with arbitrary orientations.

To obtain a realistically shaped volume conductor model for source reconstruction, despite the lack of subject-specific MR scans, we used the boundary element model for the fsaverage brain template provided by Freesurfer. Based on the volume source space and the lead-field matrix computed in the forward solution, we then computed a linearly constrained minimum variance (LCMV) beamformer [10], i.e. a spatial filter that scanned, with a set of weights, for each source location through the predefined source space grid and estimated the MEG activity at each source point independently. We thereby used a data covariance matrix estimated from a oneminute MEG data segment and a noise covariance matrix estimated from three-minute pre-stimulus empty room recordings. The beamformer was applied to the raw MEG data of each subject, leading to an estimation of a three-dimensional current dipole vector with a certain magnitude and direction at each of the source locations.

All brain plots show the two-dimensional projection of the maximum magnitude of activated voxels in the described ROI, with the ‘average’ brain template as overlay.

### Statistical Analysis

Statistical tests of the significance of neural responses on the population level were done by comparing the calculated TRFs to noise models. Noise models were created by circularly shifting the audio feature in time [43]. In detail, we circularly shifted each audio feature three times, by 15 s, 30 s and 45 s, then calculated TRFs for each of the shifted audio features and averaged across all time-shifted TRFs. This was done for each subject, resulting in two noise models (one for each audio feature) for each subject. Although the temporal relation between the audio stimulus and the MEG response is destroyed this way, the local temporal structure of the audio signal is preserved.

To assess statistical significance in the sensor-level TRFs, we bootstrapped the single-subject noise models for each audio feature. We resampled the noise models across time lags, averaged across subjects and channels and computed the magnitudes across time lags in the same way as it was done for the actual TRFs. By repeating this 10, 000 times, we obtained a distribution of noise model magnitudes across time lags. Subsequently, we estimated the relative amount of values from the noise distribution that exceeded the actual TRF for each model to therefrom estimate empirical *p*-values for each time lag. The *p*-values were then corrected for multiple comparisons using the Bonferroni method.

The statistical testing for the source-level TRFs was done analogously. For each subject and for each of the shifted audio features, we calculated source-level TRFs with the source-reconstructed raw MEG data. We also computed the corresponding noise models and averaged them across the three latency shifts. We then applied the same bootstrap algorithm as described for the sensor-level TRFs.

To assess putative laterization of cortical activity in time regions where significant responses emerged, we applied a two-tailed Wilcoxon signed-rank test on the magnitude differences of the TRFs in these time windows between the right and the left cortical ROI.

## RESULTS

We assessed the speech-FFR through two speech features. First, as already employed in our previous EEG studies, we computed a fundamental waveform from the speech signal that, at each time instance, oscillated at *f*_0_ [24, 27]. Second, because the higher harmonics of *f*_0_ contribute to the response as well, we considered envelope modulations at *f*_0_ in the higher frequency bands, a speech feature that has previously been shown to elicit strong MEG and EEG signals [38, 43].

### Temporal aspects of the neural responses on the sensor level

As a first approach, we computed temporal response functions (TRFs) that related the two speech features, the fundamental waveform and the envelope modulation, at several temporal lags, to the MEG signals at the different sensors (Fig. 2). To assess at which latencies significant neural responses emerged, we compared, for the different latencies, the amplitude of the TRFs averaged over all MEG sensors to the amplitudes of noise models, with a Bonferroni correction for multiple comparisons (Methods). Averaging across all MEG sensors may lead to a more noisy signal than when selecting particular regions of interest. However, this conservatice approach avoids bias towards particular MEG channels and captures all measured activities.

**FIG. 2.**
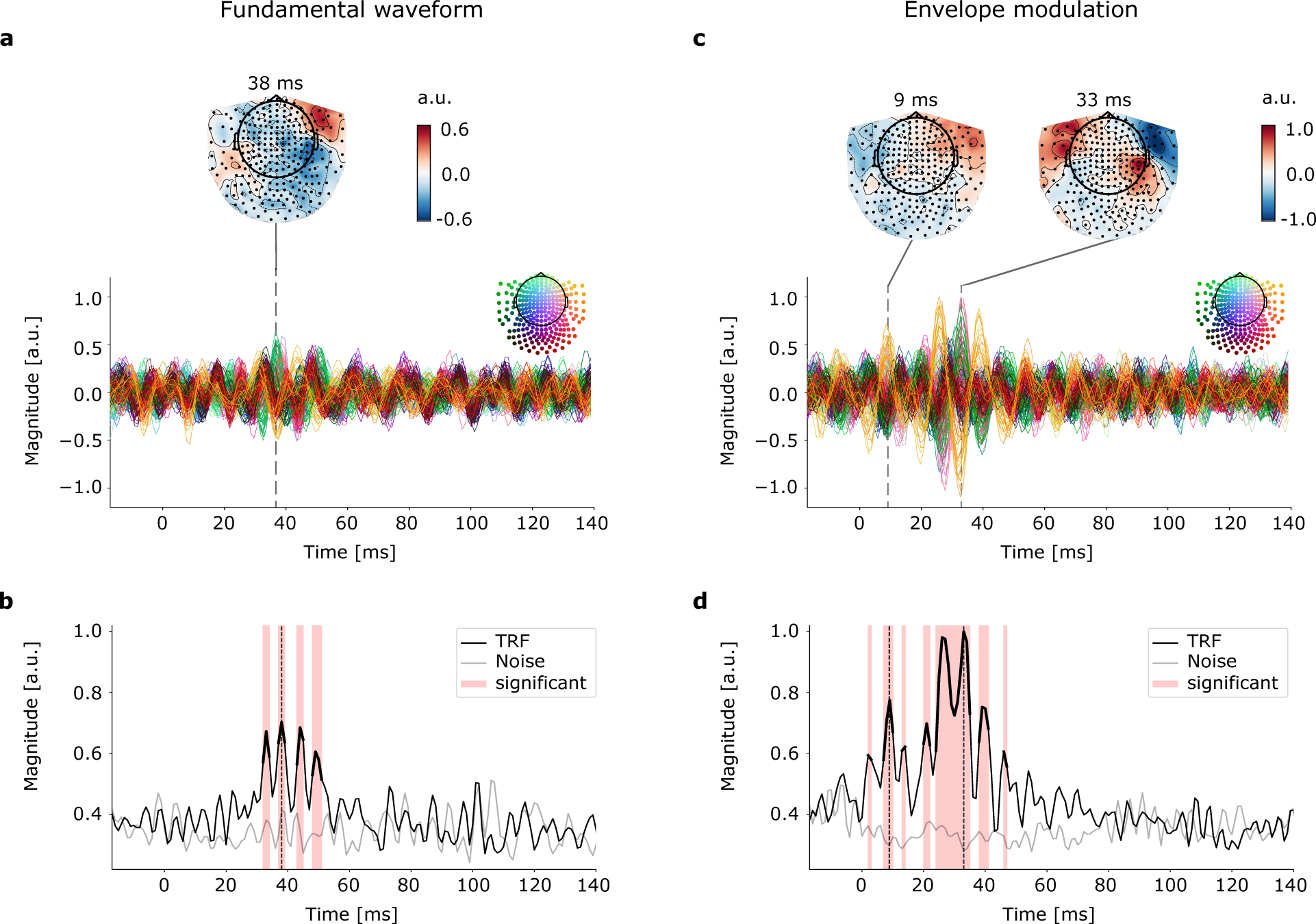
Sensor-level temporal response functions (TRFs) for the acoustic stimuli. (a), (c) The normalized sensor-level TRF for each MEG sensor for time lags between *−*20 ms and 140 ms, for the fundamental waveform (a) and the envelope modulation (c). The delays at which the amplitude of the sensor-level TRFs peak are indicated by dashed lines; the topographic plots show the corresponding sensor activations. (b), (d) The normalized absolute values of the TRFs averaged across the different sensors. The comparison of the TRF magnitudes to those of a noise model (grey line) showed that significant responses emerged around certain peak latencies (red shaded area, thicker black line, *p <* 0.05, corrected for multiple comparisons). (b) The absolute value of the TRF for the fundamental waveform displays the largest peak at a delay of 38 ms (dashed line). (d) The absolute value of the TRF for the envelope modulation peaks at delays of 9 ms and 33 ms (dashed lines).

The TRFs for the neural response to the fundamental waveform of speech yielded significant activities at time lags that ranged from 32 ms to 41 ms, with a peak activity at 38 ms (Fig. 2 (a), (b)). The topographic plot at the delay of the peak showed that the highest magnitudes occurred for MEG sensors in the right hemisphere, in particular in the right frontal and the right temporal region.

Regarding the neural response to the envelope modulation, the corresponding TRF showed significant activity between 2 - 14 ms, with peak activity at 9 ms and the highest magnitudes in the right and left frontal regions. The neural activity then peaked a second time at 33 ms, with significant activity between 20 - 47 ms, and was shaped by the left frontal, right frontal and right temporal regions. (Fig. 2 (c), (d)).

### Cortical component of the speech-FFR

In addition to the temporal characteristics, we aimed to investigate the neural origins of the measured MEG signals. We therefore performed a subject-wise source estimation on the raw, but preprocessed MEG data to localize the origin of the measured neural responses. For the cortical contribution, we determined a volume source space for a pre-defined cortical ROI (see Methods) and estimated the raw MEG signal at each voxel with a LCMV beamformer. We then computed TRFs on the source-level MEG data to relate them to both speech features at the different time lags. As already done for the sensor-level TRFs, we computed the amplitude of the TRFs, averaged over all subjects and all vertices in the cortical ROI (Fig. 3 (a)).

**FIG. 3.**
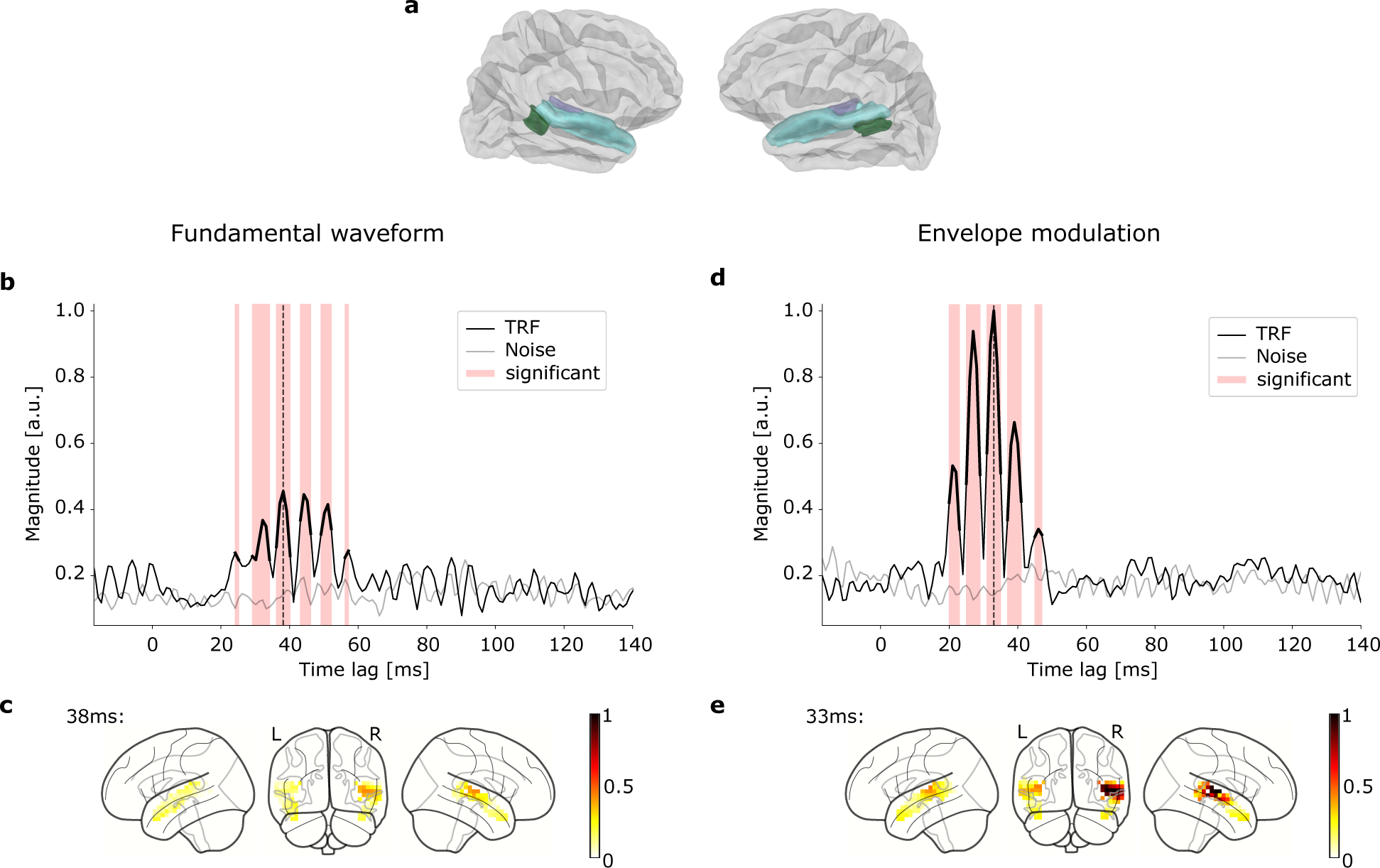
Volume source localization and source-level TRFs for the cortical ROI. (a) The cortical ROI consisted of six subregions of the Freesurfer ‘aparc+aseg’ parcellation, the right and left transverse-temporal lobe (purple), the right and left superior-temporal lobe (blue) and the tight and left banks (green). (b) The normalized amplitude of the source-level TRF for the fundamental waveform, averaged across subjects and vertices in the cortical ROI. The significant time lags (red background and thicker black line, *p <* 0.05, corrected for multiple comparisons) peak at 38 ms (dashed line). (c) The projection of the magnitudes of the voxel TRFs in the cortical ROI to the average brain template at the peak latency of 38 ms showed a dominant contribution from the right Heschl’s gyrus. (d) The normalized amplitude of the source-level TRF for the envelope modulation, averaged across subjects and vertices in the cortical ROI. The significant time lags (red) peak at 33 ms (dashed line). (e) The highest magnitudes of the voxel TRFs in the cortical ROI at the latency of 33 ms occurred again in the right transverse temporal gyrus.

The average amplitudes of the source-level TRFs were tested for significance against a noise model using a bootstrap statistic for each time lag, with Bonferroni correction for multiple comparisons. The TRF for the fundamental waveform showed significant time lags in the range of 23 ms to 57 ms, peaking at 38 ms (Fig. 3 (b)). To further analyze the origin of the peak signal at 38 ms, we projected the magnitudes of the subject-averaged voxel TRFs to the cortical ROI of the fsaverage brain template (Fig. 3 (c)). We found the highest magnitude in the right transverse-temporal part of the cortical ROI. The neural activity at this latency and in the right hemisphere was indeed significantly higher than that in the left hemisphere (Wilcoxon signed-rank test, *p* = 7 *·* 10*^−^*^5^).

Regarding the TRF for the envelope modulation, we found significant time lags in the range of 20 ms to 47 ms, peaking at 33 ms (Fig. 3 (d)). As for the fundamental waveform TRF, the average amplitudes of the source-level TRFs were tested for significance against a noise model using a bootstrap statistic for each time lag, with Bonferroni correction for multiple comparisons. The projection of the estimated amplitudes of the subject-averaged vertex TRFs on the cortical ROI of the fsaverage brain template at the peak time lag of 33 ms (Fig. 3 (d)) showed that the highest magnitude occurred again in the right-hemispheric transverse-temporal region. The TRF amplitudes obtained in the right hemisphere were indeed significantly higher than those in the left hemisphere (Wilcoxon signed-rank test, *p* = 8.8 *·* 10*^−^*^6^).

Comparing the source-level TRFs of the fundamental waveform to those of the envelope modulation, we found that the amplitude of the latter was approximately twice as large as the amplitude of the former.

In addition to the population-averaged TRFs, we also assessed subject-specific TRFs for both speech features in the cortical ROI (Fig. 4). The subject-specific TRFs were computed for each individual volunteer. They were determined to assess how reliable the neural responses could be detected at the level of individual participants, and to assess the subject-to-subject variability.

**FIG. 4.**
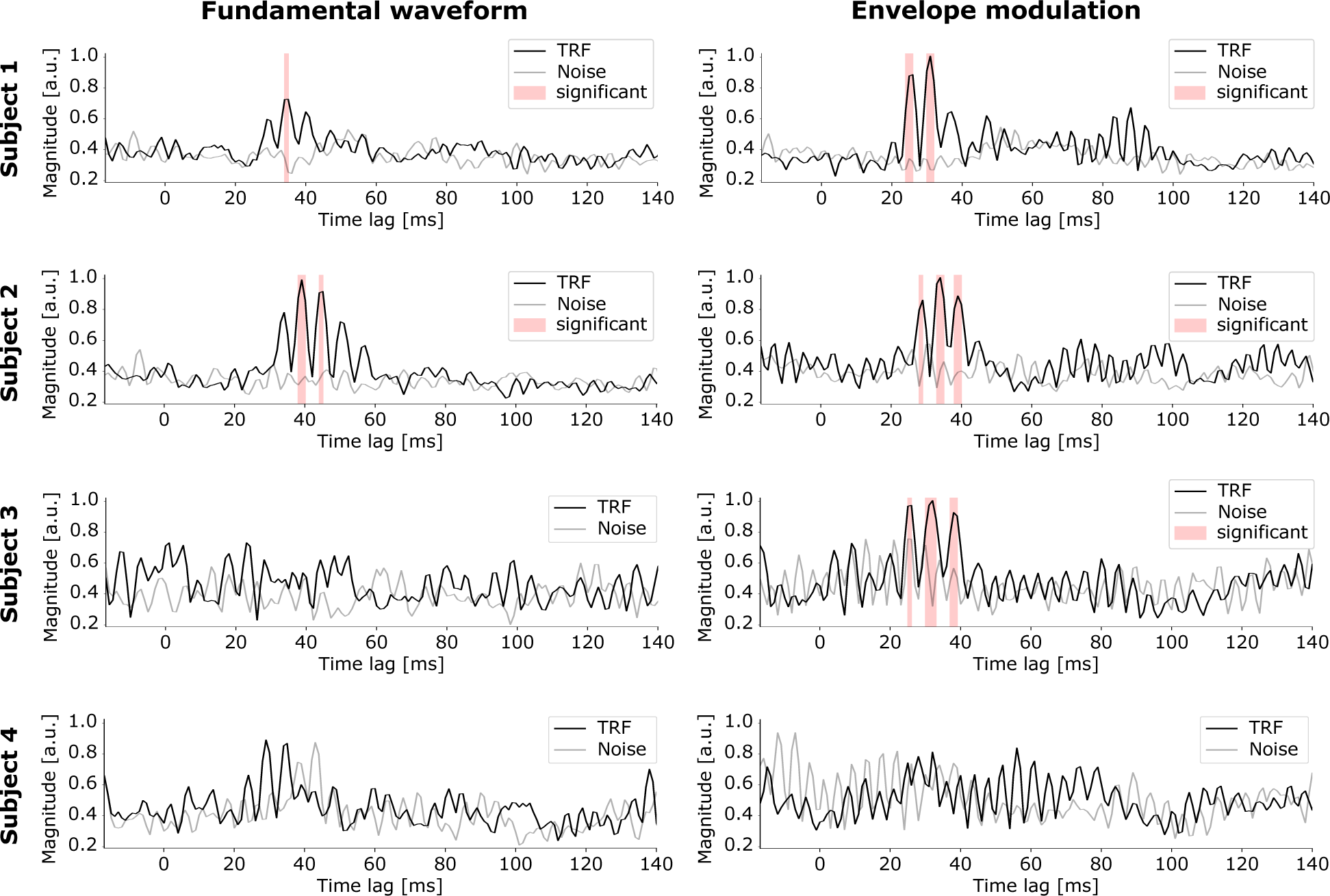
Source-level, subject-specific TRFs for the cortical ROI. The normalized amplitude of the source-level TRF for the fundamental waveform (left) and envelope modulation (right) is shown averaged across vertices in the cortical ROI for 4 subjects. Time lags at which significant neural responses occur are highlighted through a red background (*p <* 0.05, corrected for multiple comparisons). Subjects 1 and 2 are examples where significant neural responses to both speech features occur. Subjects 3 and 4, in contrast, show examples where little or no significant neural activity is detected.

As for the population-averaged TRFs, the average amplitudes of the subject-specific TRFs were tested for statistical significance against a noise model using a bootstrap statistic for each time lag, with Bonferroni correction for multiple comparisons.

The source-level, subject-specific TRFs in the cortical ROI for the envelope modulation feature showed significant peaks between 26 ms and 39 ms for 9 out of the 12 subjects. For the fundamental waveform TRFs, 4 out of 12 subjects revealed significant responses at time lags between 35 ms and 39 ms.

### Subcortical contribution to the speech-FFR

In addition to the cortical investigation, we estimated the raw MEG data on vertices of a subcortical ROI (Fig. 5 (a)), including the brainstem, to analyze possible subcortical contributions to the neural response. As for the cortical ROI, we computed TRFs on the subcortical source-level MEG data for both audio features. The average amplitudes of the source-level TRFs were again tested for significance against a noise model using a bootstrap statistic for each time lag, with Bonferroni correction for multiple comparisons.

**FIG. 5.**
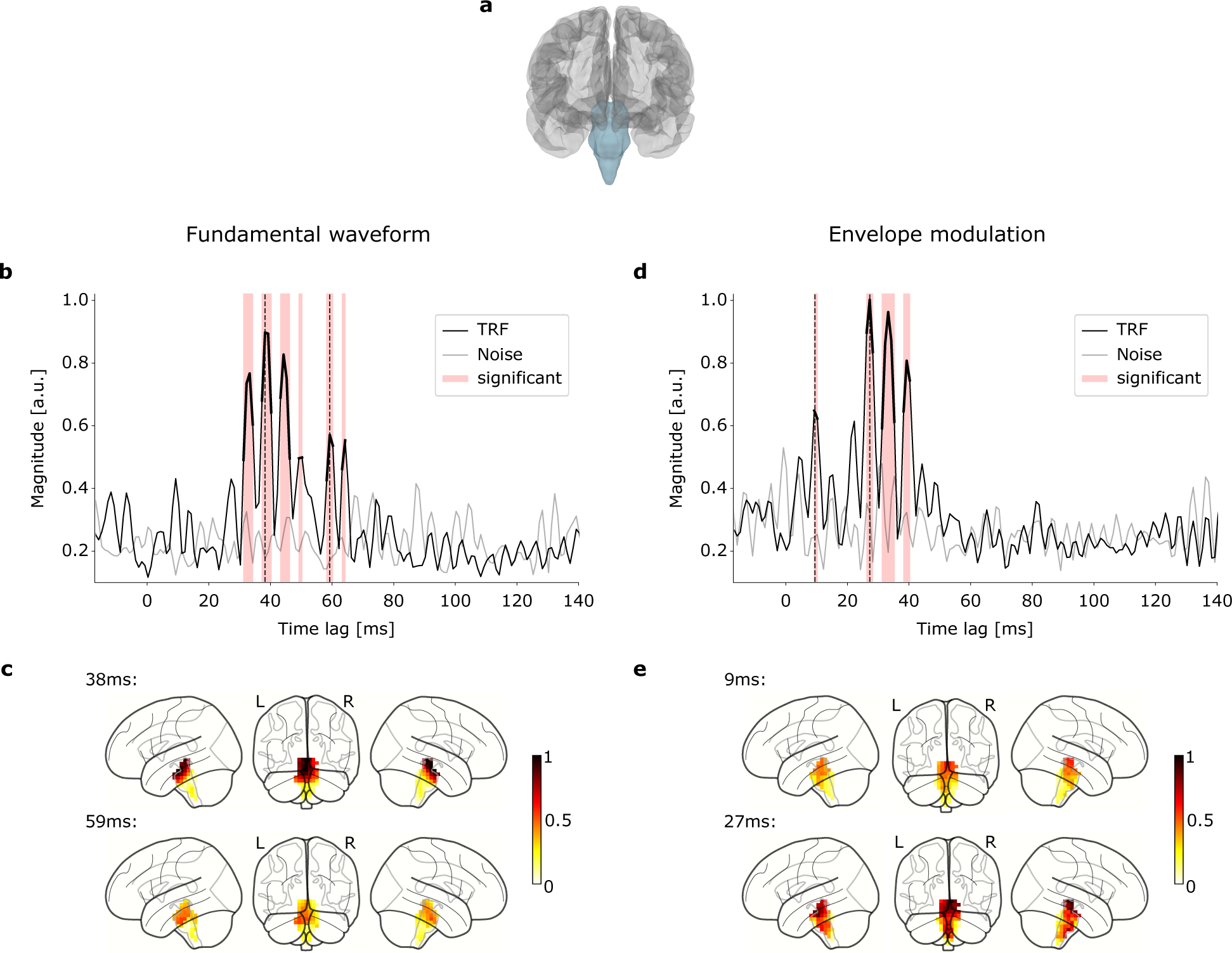
Volume source localization and source-level TRFs for the subcortical ROI. (a) The subcortical ROI consisted of the ‘brain-stem’ region of the Freesurfer ‘aparc+aseg’ parcellation. (b) The normalized amplitude of the source-level TRF for the fundamental waveform, averaged across subjects and vertices in the subcortical ROI. Significant neural responses (red background and thicker black line, *p <* 0.05, corrected for multiple comparisons) peaked at 38 ms, as well as later at 59 ms (dashed lines). (c) Projection of the magnitudes of the source-level TRFs in the subcortical ROI to the average brain template at the peak latency of 38 ms (upper panel) and at 59 ms (lower panel). (d) The normalized amplitude of the source-level TRF for the envelope modulation, averaged across subjects and vertices in the subcortical ROI. Significant responses (red background and thicker black line, *p <* 0.05, corrected for multiple comparisons) emerged around a first peak at 9 ms and around a second peak at 27 ms (dashed lines). (e) Projection of the magnitudes of the source-level TRFs in the subcortical ROI to the average brain template at the latencies of 9 ms and 27 ms.

The normalized amplitude of the subject- and voxel-averaged TRF for the fundamental waveform showed significant time lags in the range of 31 ms to 50 ms and again from 58 ms to 64 ms, peaking at 38 ms and 59 ms (Fig. 5 (b)). The projection of the magnitudes of the subject-averaged source-level TRFs at these peak latencies to the subcortical ROI, i.e. the brainstem, of the average brain template indicated highest activation in the upper brainstem region at both peak latencies (Fig. 5 (c))

For the envelope modulation feature, the normalized amplitude of the subject- and voxel-averaged TRF showed significant neural activity at time lags from 9 ms to 10 ms, peaking at 9 ms, as well as between 26 ms to 40 ms, peaking at 27 ms (Fig. 5 (d)). We projected the source-level TRF magnitudes of both peaks to the subcortical ROI of the average brain template. As it was already the case for the fundamental waveform feature, the highest activation emerged in the upper brainstem region, at both peak latencies (Fig. 5 (e)).

As for the cortical ROI, we also computed subject-specific TRFs for both speech features in the subcortical ROI (Fig. 6). The average amplitudes of the TRFs were again tested for significance against a noise model using a bootstrap statistic for each time lag, with Bonferroni correction for multiple comparisons.

**FIG. 6.**
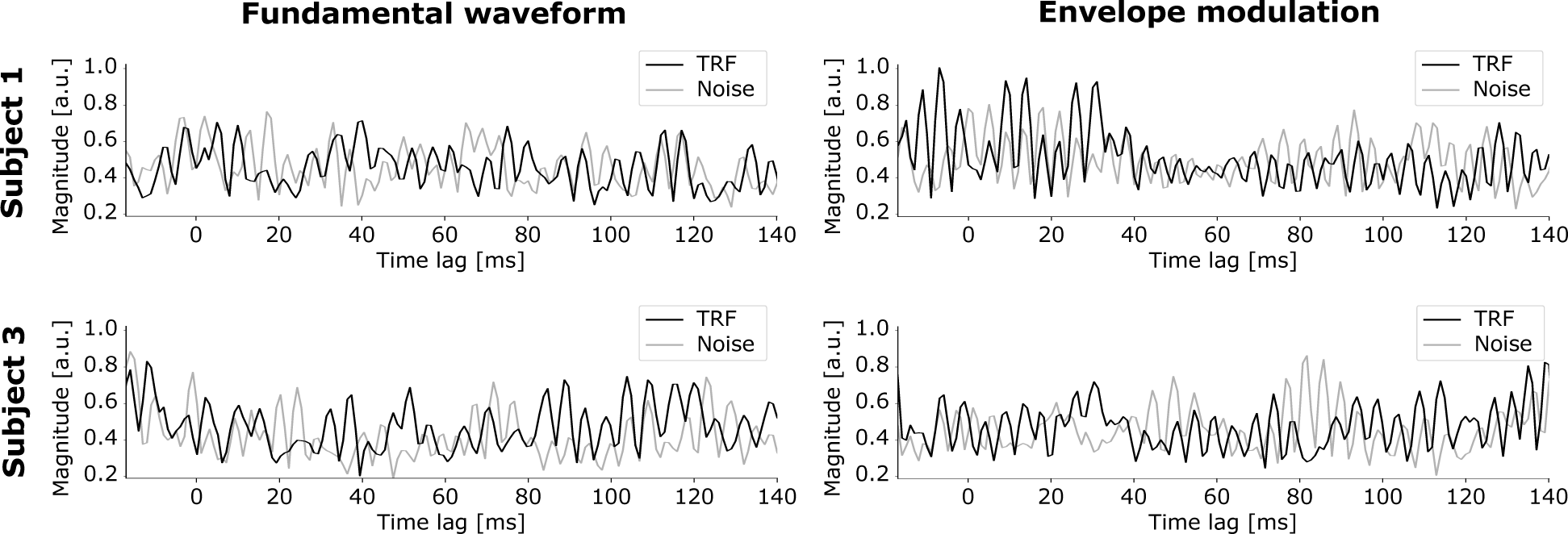
Source-level, subject-specific TRFs for the subcortical ROI, presented for two typical subjects. The normalized amplitudes of the source-level TRFs for the fundamental waveform (left) and for the envelope modulation (right) are shown averaged across vertices in the subcortical ROI. The TRF for the envelope modulation for subject 1 showed a high activity between 0 ms and 40 ms, but did not reveal any significant peak. The TRFs for subject 3 showed no significant peak for both features.

The source-level, subject-specific TRFs in the subcortical ROI for the envelope modulation as well as for the fundamental waveform showed no significant peak for any of the participants.

## DISCUSSION

In this study, we showed that the early subcortical contribution to the speech-FFR in response to continuous speech at the delay of 9 ms, previously measured with EEG in response to continuous speech, can be measured with MEG as well. We further showed that this early subcortical contribution is temporally separated from later cortical and putative subcortical activities.

### Early subcortical neural response on the population level

We considered two features of the speech stimulus: a fundamental waveform, oscillating at the speaker’s fundamental frequency of around 100 Hz, and the envelope modulation of the higher harmonics. Volume source reconstruction followed by the estimation of source-level TRFs allowed to trace the responses to these two speech features back to their neural origins in the brain. Since we performed the source reconstruction based on an average brain due to the absence of subject-specific MR scans, the interpretation of the obtained results needs to be done particularly carefully.

Importantly, we measured an early subcortical contribution to the speech-FFR. The early subcortical signal occurred in a temporal range between 7 ms and 18 ms, with a significant peak at a delay of 9 ms (Fig. 2 (d)). Importantly, the peak subcortical activity occurred much before the first cortical activity peak, at 9 ms versus 27 ms and 33 ms. The early subcortical activity was hence temporally well separated from the later cortical and putative subcortical responses.

Subcortical responses at the fundamental frequency of continuous speech had been measured before in the same range of delays through EEG, confirming that our MEG measurements relate to the same neural activity [24, 27]. EEG is known to be more sensitive to deep subcortical sources, whereas MEG is known to have more sensitivity to cortical structures, rather than to the deeper subcortical sources, due to the head tissue the signal passes. The MEG recordings in this study were obtained from a magnetometer-based MEG. This type of MEG is more sensitive to deeper subcortical structures than gradiometer-based MEG [4, 46]. Prior studies on magnetometer-based MEG responses to clicks or short speech tokens accordingly identified subcortical contributions [14, 51]. Moreover, MEG recordings found contributions from the hippocampus [17], the amygdala [18] and the thalamus [54] elicited by other types of stimuli.

Even though we see an early peak emerging in the TRF for the fundamental waveform at a delay of 9 ms (Fig. 5 (b)), this peak is not statistically significant. However, its shape resembles that of the significant early peak at the same latency in the TRF for the envelope modulation (Fig. 5 (d)), supporting the notion of the envelope modulation causing a greater neural response than the fundamental waveform, not only in cortical regions, but also regarding subcortical activity. However, this result is contrary to a recent EEG study that found a response at a delay of 10 ms tracking the speaker’s fundamental frequency and a later response at a latency of 21 ms, tracking the envelope modulation [38]. These differences might once more reflect the different neural sources measured with MEG and EEG.

### Absence of the subcortical neural response for individual subjects

The assessment of the individual subject-specific TRFs for the subcortical ROI revealed no significant responses, for neither of the two speech features (Fig. 6). This finding presumably reflects the lack of sensitivity of MEG to subcortical sources, hindering their detection on the level of individual subjects. This finding also reinforces the need of a population average to reveal the early subcortical response from MEG.

### Cortical neural response on the population level

The subject- and voxel-averaged source-level TRF for the cortical ROI in response to the fundamental waveform exhibited at the peak latency time of 38 ms (Fig. 3 (b)) and revealed a dominant cortical origin in the transverse-temporal region. The source-localized cortical activity in response to the envelope modulation peaked at a latency of 33 ms. It had a greater amplitude than the response to the fundamental waveform, matching our findings in the sensor-level TRFs that suggested the envelope modulation feature to have a greater contribution to the neural response than the fundamental waveform. These results are also consistent with a recent MEG study that found a cortical response to the envelope modulation of the higher harmonics of continuous speech, emerging at a latency of about 40 ms [43].

Despite the limitations posed by the use of an average MR scan and a 5 mm source grid, our results show the presence of cortical responses at the fundamental frequency between 20 ms and 57 ms. Although natural continuous speech is much more complex than the repeated syllables used in a previous study on these neural responses, our observations match the long-lasting explanatory power of the neural responses in the auditory cortex described there [14].

The neural responses in the cortical ROI to both speech features showed the highest magnitude in the transverse-temporal gyrus (Fig. 3 (c) and Fig. 5 (e)), also known as Heschl’s gyrus, which defines the primary auditory cortex region. This is in line with previous studies indicating latencies around 40 ms to occur from Heschl’s gyrus [9, 45, 61].

We found a significant right laterization in the cortical ROI for the neural responses to both the fundamental frequency and the envelope modulation feature. This phenomenon has already been shown in previous MEG studies on continuous speech [43] as well as for short speech tokens [14]. Moreover, prior studies on neurophysiological processing of voice information with functional magnetic resonance imaging (fMRI) showed the right hemisphere to play a fundamental role in spoken language comprehension [44]. Further studies on fMRI responses observed the right laterization using sung speech stimuli, indicating the presence of a brain asymmetry for speech and melody [2]. Motivated from prior studies that showed the right auditory cortex to be specialized for early tonal processing and pitch resolution, it has been suggested that the cortical responses occur as a consequence of early auditory processing of acoustic periodicity [12, 37, 43].

### Cortical neural response for individual subjects

Assessing individual subject-specific TRFs allowed us to investigate the variability in the speech-FFR across individuals. Our findings demonstrate that the speech-FFR evoked by continuous speech in the cortical ROI can not only be detected at the population level but also at the level of individual participants, providing further evidence for the robustness of this response.

In particular, we found that the neural response to the envelope modulation showed significant peaks in the source-level, subject-wise TRFs in the cortical ROI (Fig. 4) for the majority of the participants (9 out of 12).

In contrast, the TRFs for the fundamental waveform showed significant peaks in only a subset of participants (4 out of 12), underlining the prior findings that the envelope modulation leads to a greater neural signal than the fundamental waveform [43].

### Late subcortical responses

The subject- and voxel-averaged source-level TRFs yielded significant subcortical activity around the later time lags of 27 ms, 38 ms, and 59 ms (Fig. 5). These neural activities were mostly visible in the upper brainstem and less in the deeper brainstem structures. Because of the comparatively late timing, matching that of the stronger cortical responses, these neural activities may in fact stem from dispersed cortical activity and may have appeared in subcortical sources due to volume conduction and associated lubrication effects.

Volume conduction happens when electrical activity propagates from one part of the brain to another through the surrounding tissues and fluids, which may result in the appearance of activity in brain regions where it does not actually occur [50, 52]. Lubrication effects refer to the smoothing of source estimates in the brain due to the use of an average brain template for source reconstruction, which can also result in activity appearing in other brain regions [5, 19, 31]. Both types of effects are indeed common and cannot be neglected when interpreting source reconstruction results that employ, in the lack of individual structural MRIs of the participants, an average brain MRI.

### Differences in neural responses to the fundamental waveform and to the envelope modulation

In this study, we used two speech features, the fundamental waveform and the envelope modulation, to investigate the speech-FFR elicited by continuous speech. Using temporal response functions (TRFs), we examined how neural responses to these different stimulus features arise in subcortical as well as cortical areas. Our results showed that both the envelope modulation and the fundamental waveform drive significant cortical responses. However, the envelope modulation caused a larger neural response than the fundamental waveform.

Regarding subcortical activity, we only observed significant activity in response to the envelope modulation, and not in response to the fundamental waveform. The latter was presumably too small to be detected, reinforcing the notion that the envelope modulation drives a stronger neural response than the fundamental waveform, both on the subcortical and on the cortical level.

Previous studies have indeed shown the perceptual relevance of envelope modulation in speech understanding, particularly above 300 Hz, as well as its greater resistance to background noise as compared to the lower frequencies below 200 Hz [3]. This perceptual relevance might be reflected in the larger neural response to the envelope modulation as opposed to the fundamental waveform that we observed here.

Moreover, previous studies on FFRs to specially-designed tones discovered that the response at the fundamental frequency emerges even when that frequency itself is missing from the stimulus, as long as higher harmonics are present [28, 56]. The extensive nonlinearities in the auditory system, starting from the compressive nonlinearity in the inner ear and continuing through the nonlinearities associated to neural responses, can indeed extract the fundamental frequency from the higher harmonics. This mechanism appears to dominate over the direct neural response to the fundamental frequency itself.

In line with our findings, a previous MEG study on the speech-FFR elicited by continuous speech likewise obtained larger cortical responses to the envelope modulation as compared to the fundamental waveform [43]. In addition, one of our earlier EEG investigations into this issue found that the envelope modulation explained a larger variance of the neural data than the fundamental waveform, further supporting our findings on the subcortical level [38]. Taken together, these previous findings as well as our current ones suggest that the envelope modulation is the more important speech feature for assessing the speech-FFR to continuous speech than the fundamental waveform.

## CONCLUSIONS

In summary, we simultaneously recorded early subcortical and cortical contributions to the speech-FFR using natural continuous speech and MEG. Using TRF analysis and neural source estimation, we showed that it is possible to separate contributions to the neural response from cortical and subcortical sources and moreover to assign those signals to two different features of continuous speech. This provides the opportunity of TRF analysis applied on MEG recordings to investigate further research questions on auditory processing of continuous speech under several stimulus conditions. The temporal separation of cortical and subcortical neural signals may allow to investigate the involvement of early subcortical responses in higher cognitive aspects of speech processing, such as attention to one of several competing speakers. Moreover, the powerful properties of MEG such as the fine temporal resolution, as well as the high spatial resolution, may allow an improved investigation of interactions between subcortical and cortical structures during auditory processing.

## DATA AVAILABILITY

The MEG data used in this study is available from the authors upon request.

## AUTHOR CONTRIBUTIONS

**Alina Schüller**: Conceptualization, Interpretation of Results of Experiments, Formal analysis, Investigation, Writing original draft. **Achim Schilling**: Data Curation, Interpretation of Results of Experiments, Reviewing and Editing of the paper. **Patrick Krauss**: Data Curation, Interpretation of Results of Experiments, Reviewing and Editing of the paper. **Tobias Reichenbach**: Conceptualization, Interpretation of Results of Experiments, Investigation, Supervision, Reviewing and Editing of the paper.

## ACKNOWLEDGMENTS

The authors are grateful to the publishers Deuticke Verlag and Hörbuch Hamburg for the permission to use the novel and corresponding audio book *Gut gegen Nordwind* by Daniel Glattauer for the present and future studies.

This work was funded by the Deutsche Forschungsgemeinschaft (DFG, German Research Foundation): grant KR 5148/2-1 to PK (project number 436456810) and grant SCHI 1482/3-1 (project number 451810794) to AS, and by the Emerging Talents Initiative (ETI) of the University Erlangen-Nuremberg (grant 2019/2-Phil-01 to PK).

